# 16s rRNA gene sequence analysis of the microbial community on microplastic samples from the North Atlantic and Great Pacific Garbage Patches

**DOI:** 10.1101/2022.12.22.521553

**Authors:** Dkawlma Tora, Ute Hentschel, Stefan Lips, Mechthild Schmitt-Jansen, Erik Borchert

## Abstract

The exponential increase in plastic production has led to their accumulation in the environment, particularly in oceans, polluting these environments from the shore to the open ocean and even sea ice in the pole regions. We compared microbial communities on plastic particles, known as “Plastisphere”, collected from the Atlantic and Pacific oceans gyres in the Summer of 2019 and subsequently looked for potential plastic degraders. We applied a 16S rRNA amplicon sequencing approach to decipher differences and similarities in colonization behaviour between these two gyres. Two polymer types include plastics: polyethylene (PE) and polypropylene (PP). We found that microbes differed significantly between the two oceans and identified thirty-two differentially abundant taxa at the class level. Proteobacteria, Cyanobacteria and Bacteroidota were the most prominent relative abundant phyla in the two oceans. Finally, according to the current literature, we found 40 genera documented as potential plastic degraders. This study highlights the importance of the biogeographical location with respect to microbial colonization patterns of marine plastic debris, differing even in the open oceans. Furthermore, the wide distribution of potential plastic-degrading bacteria was shown.

## INTRODUCTION

4.8 to 12.7 million metric tons of plastics are estimated to enter the ocean yearly, mostly from land (Jambeck et al., 2015). Dris et al. (2016) related an atmospheric fallout between 2 and 355 particles/m^2^/day. Microscopic plastic fragments and fibres (microplastics) are widespread in the oceans. They have accumulated in the pelagic zone and sediments resulting from the degradation of macroplastic items (Thompson et al., 2004). The ubiquitous plastics in the ocean could harm the marine environment and humans through the food web. Evidence showed that microplastics could act as passive samplers for toxic compounds such as persistent organic pollutants (e.g., industrial chemicals, dioxins, pesticides) and heavy metals from seawater, leading to an increased negative impact on the biota (Mato et al., 2001; Horn et al., 2019). Besides that, the potential accumulation of microplastics in the food chain, especially in fish and shellfish, also exposes human consumers to these adsorbed chemicals (Kershaw & Rochman, 2014). Lusher et al. (2013) reported that 36% of pelagic and demersal fish collected from the English Channel had microplastics in their gastrointestinal tract.

Outreaches, national and international laws, policies, and conventions have been discouraging the use of plastic and its release into the environment to face plastic pollution. The African continent is at the forefront of legislative actions against plastic pollution. For instance, Rwanda has banned non-biodegradable plastic since 2008 and single-use plastics in 2019. The ban prohibited the manufacturing, use, import and sale of plastic carrier bags and forbade travellers into Rwanda to come with such products. Nigeria announced a ban on plastic bags in 2013, and in 2020, it strengthened its legislation by including a fine of 1072.16 Euro or three years jail term for any store found giving plastic bags to customers. In Botswana, a minimum thickness for bags was established and mandated that retailers apply a minimum levy to thicker bags, which would support government environmental projects. Kenya has the strictest ban on single-use plastic globally and in protected areas (Greenpeace, 2021).

Plastic is a high molecular weight synthetic polymer of a long chain of hydrocarbons derived from petrochemicals (Ahmed et al., 2018). With swift development in molecular techniques, research focused on microbial communities living on plastics and their ability to degrade hydrocarbons. The biological deterioration of plastic pollutants depends on many factors: surface area, functional groups, molecular weight, hydrophilic and hydrophobicity, melting temperature, chemical structure, crystallinity, etc. (Okada, 2002). Microbial degradation of plastic involves many steps: biodeterioration, bio-fragmentation, assimilation, and mineralization (Purohit et al., 2020).

Zettler, Mincer & Amaral-Zettler (2013) coined the term “Plastisphere” to describe biofilm-forming communities on marine plastic debris. They collected marine plastic debris at multiple locations in the North Atlantic to analyze the microbial consortia attached to it. They found diverse microbial communities, including heterotrophs, autotrophs, predators, and symbionts, which they called a ‘Plastisphere’. Coons et al. (2021) investigated plastic-type and incubation locations in the Atlantic and Pacific oceans, focusing on shore locations as drivers of marine bacterial community structure development on plastic via 16S rRNA gene amplicon analysis. They found that incubation location was the primary driver of the coastal Plastisphere composition. The bacterial communities were consistently dominated by the classes Alphaproteobacteria, Gammaproteobacteria, and Bacteroidia, irrespective of sampling location or substrate type.

Similarly, in 2015, Amaral-Zettler et al. used next-generation DNA (Deoxyribonucleic Acid) sequencing to characterize bacterial communities from the Pacific and Atlantic oceans. Their objective was to determine whether the composition of Plastisphere communities reflects their biogeographic origins. They found that these communities differed between ocean basins and, to a lesser extent, between polymer types and displayed latitudinal gradients in species richness.

For this work, we collected plastic particles from the North Atlantic and the Great Pacific Garbage Patches in 2019 and compared microbial communities from the Atlantic to the Pacific, as well as looking for potential plastic degraders. The North Atlantic and the Great Pacific Garbage Patches are the biggest current patches, with a density of 10^6^ km^-2^ (Eriksen et al., 2014) and 96 400 million metric tons of plastic (Ritchie & Roser, 2018), respectively. In addition to the work of Zettler et al. (2015), we pointed out communities responsible for the discrepancy. Besides, our samples were composed of PP and PE polymers for which only scarce information regarding environmental degradation is available. To reach our goal, 16S rRNA gene amplicon sequence analysis was used on the microbial community of these microplastic samples.

## MATERIALS AND METHODS

### Plastics collection

The samples were collected in the North Atlantic and Great Pacific Garbage Patches. The pieces from the Atlantic were collected between 26-08-2019 and 04-09-2019 during the POS536 cruise project ‘Distribution of Plastics in the North Atlantic Garbage Patch’ (DIPLANOAGAP) aboard the German research vessel (R/V) Poseidon. A Neuston catamaran onboard R/V Poseidon, equipped with a microplastic trawl net (mesh size 300 μm, mouth opening 70 cm x 40 cm) was used to collect the plastic samples from the sea surface. After each tow, all microplastic fragments were removed from the trawl sample and conserved in a saturated ammonium sulphate solution (700 g/l ammonium sulfate, 20 mM sodium citrate, 25mM EDTA, pH 5.2). This solution precipitated all proteins, preventing DNA and RNA degradation for an extended time, even at room temperature. Verification of plastic type by Attenuated Total Reflectance Fourier-Transform Infrared (ATR-FTIR) spectroscopy analysis was subsequently performed by TUTECH GmbH in Hamburg, Germany.

Another cruise project, MICRO-FATE, aboard another German R/V, the Sonne (SO268/3), between 05-06-2019 and 27-06-2019, was used to collect plastic samples at the sea surface in the Great Pacific garbage patch. Plastics were collected using a scoop net sampling method. The plastic surfaces were scraped using a flame-sterilized scalpel, and biofilms were transferred into microcentrifuge tubes. The sampling area was 16 x 16 mm, and tubes were immediately frozen in liquid nitrogen. At each station, 1 litre of pacific water was filtered through a 3 μm filter (3 μm Isopore TSTP 04700 Millipore, Merck KGaA, Frankfurt, Germany) and a 0.22 μm filter (0.22 μm Isopore GTTP04700 membrane filters Millipore, Merck KGaA, Frankfurt, Germany). Also, the filters were transferred to microcentrifuge tubes and immediately frozen in liquid nitrogen.

### Extraction of nucleic acids from the samples

For Atlantic samples, sections of the different plastic samples were cut with a sterile scalpel and placed into 2 ml MP Biomedicals™ Lysing Matrix E tubes (MP Biomedicals, Eschwege, Germany). Then physically disrupted using a bead-beating technique, with a single cycle of 30s at a speed of 5.500 rpm in a FastPrep homogenizer (Qiagen, Hilden, Germany). The DNA extraction from the lysis product was then performed using the Qiagen AllPrep DNA/RNA Minikit according to the manufacturer’s instructions. The quality and quantity of the DNA extraction were assessed using a NanoDrop Spectrophotometer (Desjardins & Conklin, 2010). The 16S rRNA gene was amplified with the primer pair 27F and 1492R. Then, the quality was checked through polymerase chain reaction (PCR). The sequencing of the V3-V4 region of the 16S rRNA gene was performed with v3 chemistry on a MiSeq Illumina sequencing platform at the Competence Centre for Genomic Analysis (CCGA) Kiel, Germany after the PCR products were visually assessed using 1% gel electrophoresis. For amplicon sequencing, the amplification of the V3-V4 hypervariable region of the 16S rRNA gene was accomplished using primer pair 341F (50-CCTACGGGAGGCAGCAG-30; Muyzer et al., 1993) and 806R (50-GGACTACHVGGGTWTCTAAT-30; Caporaso et al., 2011). Raw reads were archived in NCBI under the BioProject number PRJNA901861.

For Pacific samples, DNA was extracted from the biofilm pellets and water filters using the Macherey Nagel DNA Nucleo spin soil kit (Nucleo Spin TM Soil kit Macherey-Nagel TM, Düren, Germany) according to the manufacturer’s instructions. DNA concentration was measured using a nano Qubit (ThermoFisher). Next-generation Illumina Sequencing was performed on an Illumina MiSeq platform using a V3 (300bp paired-end read) kit with a sequencing amount of 20 million reads, using the 341F (CCTACGGGNGGCWGCAG) and 785R primer set (GACTACHVGGGTATCTAAKCC). Raw reads were archived in NCBI under the BioProject number PRJNA837054.

### Quantitative Insights Into Microbial Ecology (QIIME2) pipeline

The Raw amplicon sequences were then processed using the open-source Quantitative Insights into Microbial Ecology (QIIME2, version 2020.11) following a pipeline developed by Kathrin Busch (Busch et al., 2021). In brief, the *cutadapt* plugin was used to trim forward primers, heterogeneity spacers from forward-only single-end fastq files (Martin, 2011) and the *qualityfilter* plugin (Bokulich et al., 2013) was used to check the quality of the demultiplexed reads. An interactive plot served to visualize these results and to determine an appropriate truncation length. Then, the reads were truncated through the DADA2 algorithm to produce a total read length of 270 nucleotides. That truncation significantly increased the quality of the reads, reduced the overlap between forward and reverse reads, and allowed us to use only forwards reads for the analysis. Before the truncation, the reads were denoised using the *denoise-single* method of the DADA2 algorithm (Callahan et al., 2016), which removed chimeric sequences and inferred sample composition using a parametric error model.

The amplicon sequence variants (ASV; Callahan et al., 2017) were classified at 80% confidence level using the most recent SILVA 138 16S rRNA gene reference database (Quast et al., 2013; Yilmaz et al., 2014). Common eukaryotic contaminants (chloroplasts, mitochondria) and unassigned sequences were removed using the *filter-features* method of the *featuretable* plugin, and the resulting dataset was rarefied to 8,000 sequences. Alpha rarefaction curves have an excellent saturation for 8,000 sequences. A phylogenetic backbone tree was built using FastTree (Price et al., 2009; Price et al., 2010) and MAFFT (Katoh & Standley, 2013) alignment through the *phylogeny* plugin. The resulting tree was used to compute core diversity metrics which served to compute downstream analyses along with an alpha-rarefaction curve *via* the *diversity* plugin.

### Alpha and Beta diversity measures

The alpha diversity was investigated according to unique ASVs per sample (species richness), taking into consideration the number of times each ASV occurs in the sample (Pielou’s evenness) and the phylogenetic relatedness of each sample community (Faith’s PD). ‘*Qiime diversity alpha-group-significance*’ plugin in QIIME2 was used to assess the diversity within each area. The results were displayed through Kruskal-Wallis (all groups) and Kruskal-Wallis (pairwise) results.

Non-phylogenetic (evenness) and phylogenetic (Faith’s PD) diversity indices were visualized using the online tool QIIME2 view (https://view.qiime2.org/). Eventually, if the comparison revealed a significant difference in microbial diversity, Kruskal-Wallis pairwise was considered among groups to see where the difference lies.

Beta diversity measures assessed the differences between groups following the different parameters. ‘Qiime diversity beta-group-significance’ plugin in QIIME2 was used for this analysis. The analysis was performed using the non-metric multidimensional scaling method (NMDS; Kruskal, 1964) with a sample-wise unweighted UniFraq distance matrix (Lozupone & Knight, 2005). Each group was assessed based on its distance from the other groups in QIIME2; boxplots were displayed simultaneously with the PERMANOVA results and pairwise PERMANOVA results between groups. The PERMANOVA group significance and pairwise tests were run simultaneously through the *betagroup-significance* method (non-parametric MANOVA; Anderson, 2001) of the diversity plugin with an unweighted UniFraq matrix and 999 permutations as input.

We adopted the standard significant measure, p-value = 0.05, for these statistical analyses. All the p-values below this standard describe a significant difference between the compared parameters and vice versa.

### Different taxonomic level analysis

The feature ASVs table was exported in biom format in QIIME2. Subsequently, the taxonomy metadata file was added to the biom file and exported in TSV file format using ‘*biom convert*’ plugin in QIIME2. Further analyses outside the QIIME2 environment, such as the share of ASVs between the samples, were performed using the resulting TSV file table. Besides that, the same feature table was collapsed at the genus level (to perform the sunburst plots, which helped to display microbial communities on plastics) and the class level (to plot the differentially abundant taxa) using the ‘*qiime taxa collapse*’ plugin. The Linear discriminant analysis (LDA) effect size (LEfSe) helped to plot the differentially abundant classes between the Atlantic and the Pacific Plastisphere, utilizing ‘galaxy online’ (https://huttenhower.sph.harvard.edu/galaxy/). The level 3 data was used, arranged within Excel (according to the different oceans) and imported into Galaxy for LEfSe analysis. The analyses were performed on the microbial community relative abundance data in both oceans. Grouped data were first analyzed using the Kruskal-Wallis test with a significance threshold of 0.05 to determine if the data was differentially distributed between groups.

## RESULTS & DISCUSSION

Samples collected comprised 68 microplastic pieces from the North Atlantic and the Great Pacific Garbage Patches as well as 14 water samples from the Great Pacific Garbage Patch. The North Atlantic Garbage Patch accounted for 30 plastic samples composed of 25 PE (polyethylene) and 5 PP (Polypropylene) particles, according to FTIR analysis. In contrast, the Great Pacific Garbage Patch accounted for 38 plastic samples, composed of 28 PE and 10 PP (Supplementary table S1).

Processing all samples in QIIME2 yielded 11,852 demultiplexed ASVs. Pacific plastic accounted for 7,081 ASVs showing more richness than Atlantic plastic which accounted for 4,454 ASVs and, in turn, showed more richness than Pacific water (3,623 ASVs). The Pacific plastisphere displayed the highest number of taxa at most taxonomic levels except the phylum level (see Table 1). Here, the Atlantic plastisphere displayed the highest number of phyla with 35, whereas Pacific plastic contained 33 phyla and Pacific water 27 phyla. Overall, the Pacific water displayed the lowest amount of taxa irrespective of the taxonomic level, which might hint towards a microhabitat formation on the plastic particles as they travel across the oceans and enrich their community along the way. These results also show an increasing proportion of unclassified taxa as one moves from the phylum level to the genus level, which informs that much is still to be discovered. The Shannon diversity indice values are between 4.88 and 8.75 in the individual samples (Supplementary table S1), with no apparent large differences between the different sample types and locations.

**Table 1.**
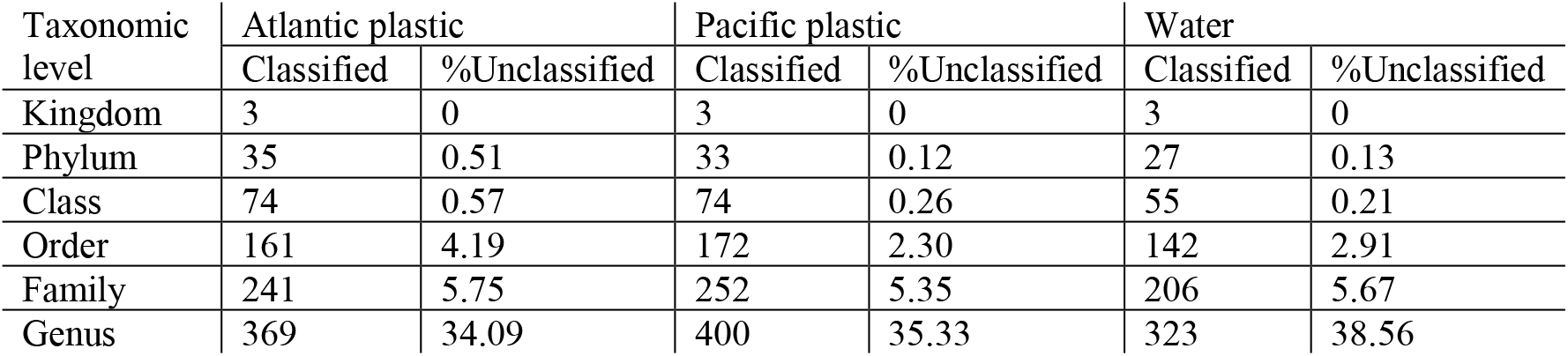
Taxonomic rank abundance distribution per sample type and percental display of unclassified ASVs per taxonomic rank.

### The Principal Coordinates Analysis (PCoA) of all samples

A PCoA plot, grouping all the samples, was generated from QIIME2 to infer the phylogenetic relatedness between the samples’ communities of both oceans. Three clusters, as shown in Figure 1, were formed. It shows that the communities of each area are related with no interference with the communities of the other location. However, in the Pacific Ocean, two clusters were formed, which could explain that an occurrent factor influences the diversity of the microbes. The outgroup in the Pacific Ocean is the only sample with a noticeable amount of Archaea belonging to the class of Thermoplasmata (0.6% of the reads). These Archaea were investigated by Gupta et al., 2021, and were shown as acidophiles.

**Figure 1.**
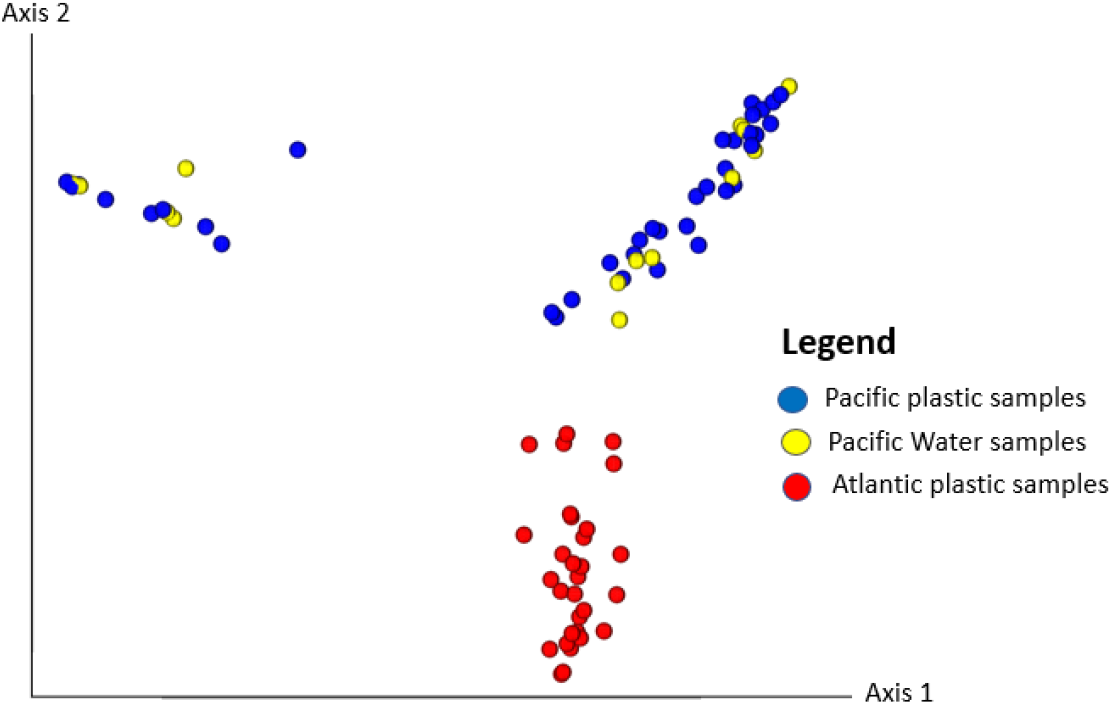
Principal coordinates analysis (PCoA) of all samples showing a specific clustering pattern of the associated microbial communities from the different oceans and samples. The phylogenetic distances calculated within the dataset indicate three clusters showing their level of relatedness. The plot was generated in QIIME2.

### The ASVs distributions between the samples

The three different sampling domains shared 611 ASVs representing 5% of the total reads. However, 380 ASVs representing 3%, were exclusively shared between Atlantic plastic and Pacific plastic, 106 (1%) between Atlantic plastisphere and Pacific water, and 1598 (13%) between Pacific Plastisphere and Pacific water. Conversely, 4492 (38%) of the ASVs were unique to the Pacific plastics, 3357 (29%) to the Atlantic plastics and 1308 (11%) to the Pacific water (Figure 2). A negligible proportion of ASVs is shared between the two oceans, while each ocean showed a big proportion of unique ASVs, suggesting a profound difference between their communities.

**Figure 2.**
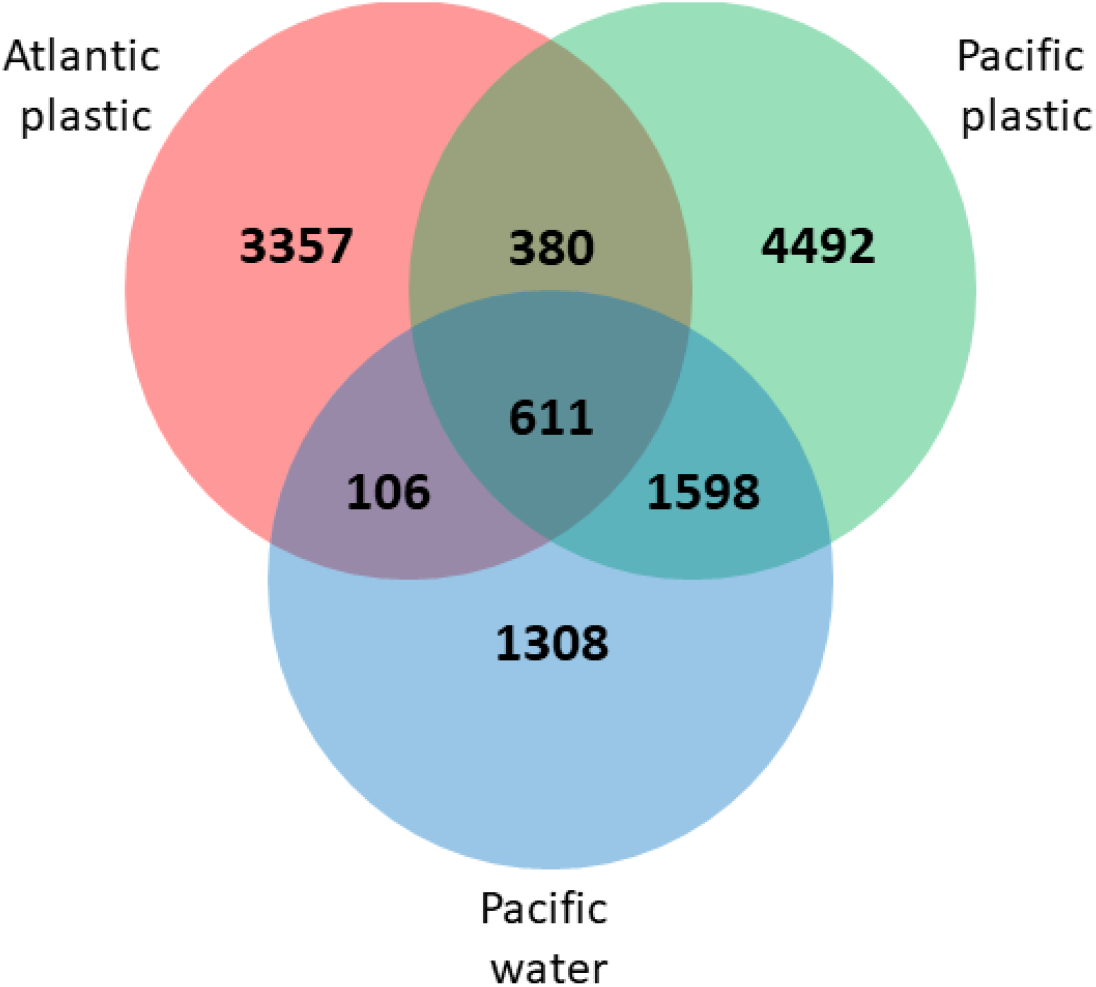
Distribution of ASVs between the different sample types. Unique and shared ASVs between Atlantic plastic, Pacific plastic, and Pacific water samples. The plot was made in R using the ‘Venn diagram’ package.

### Microbial composition on the Atlantic plastics

From the analysis, the highest relative abundances were bacteria (99.91%). Three bacterial phyla accounted for more than 90% of the relative abundance. Verrucomicrobiota, Bdellovibrionota and Firmicutes accounted for more than 1% each, while 29 other phyla (including bacterial, archaeal and eukaryal phyla) accounted for 4.70% of the community (each of these 29 phyla accounted for below 1% of the relative abundance).

Among the abundant minor domains, Eukaryota (0.09%) were represented by the phyla Amorphea (0.08%) and SAR (0.002%) and the classes of Obazoa and Alveolata. Likewise, the reads of Archaea (0.0002%) were represented by the phylum of Nanoarchaeota and the class of Nanoarchaeia.

Proteobacteria, Cyanobacteria and Bacteroidota were the three most abundant groups at the phylum level (Figure 3). The occurrent communities include Alphaproteobacteria (34.60%), reported as early colonizers; Bacteroidia (17.04%), reported as secondary colonizers and Gammaproteobacteria (10.9%), later-stage colonizers at the class level, according to a recent 16S rRNA gene amplicon data meta-analysis from 35 Plastisphere studies, which revealed the successive colonization of the Plastisphere (Wright et al., 2020). So, Gammaproteobacteria’s presence suggests the maturity of the biofilm, indicating that the plastics have been drifting for quite some time. Meanwhile, members of the phylum Cyanobacteria; have been reported as abundant components of plastic debris communities (Salta et al., 2013) highly represented on PP and PE items (Zettler et al., 2013).

**Figure 3.**
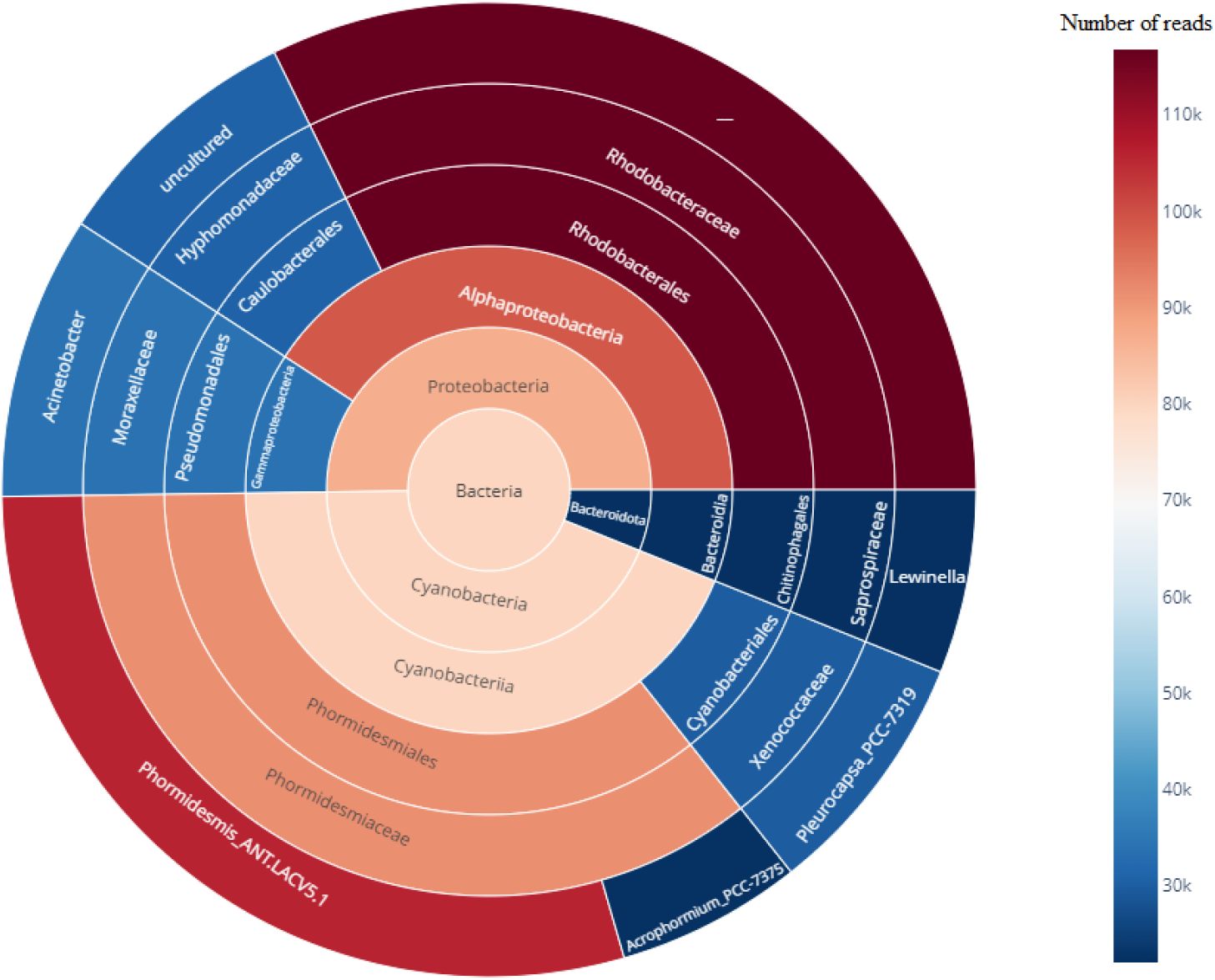
Reads and taxonomic affiliation of recurring communities on the Atlantic Plastisphere. Sunburst chart displaying the affiliations of genera that reached values above 20,000 reads. Each plot crown represents one taxonomic level from the Kingdom to the genus. The plot was implemented using ‘plotly.express’ package in Python.

Other communities at the Family level include bacteria that prefer a surface-attached lifestyle, such as Saprospiraceae (McIlroy & Nielsen, 2014), Hyphomonodaceae, known to be biofilm formers (Abraham & Rohde, 2014) and Rhodobacteriaceae as opportunistic colonizers (Dang & Lovell, 2016). At the genus level, Lewinella and Acinetobacter were described as potential plastic degraders (see Table 3).

**Table 2.** Statistical analysis of the samples: table displaying p-values from Kruskal-Wallis and PERMANOVA analysis.

|  |  | Atlantic<br>Plastisphere | Pacific<br>Plastisphere | Pacific plastics &<br>water | Atlantic & Pacific<br>plastics |
| --- | --- | --- | --- | --- | --- |
| Alpha<br>diversity | Non-phylogenetic measure |  |  |  |  |
|  | Considering the polymer<br>types | 0.8 | 0.6 | 0.96 | 0.82 |
|  | Regardless of the polymer<br>types |  |  | 0.88 | 0.38 |
|  | Phylogenetic measure |  |  |  |  |
|  | Considering the polymer<br>types | 0.67 | 0.57 | 0.4 | 0.002 |
|  | Regardless of the polymer<br>types |  |  | 0.23 | 0.000063 |
| Beta<br>diversity | Considering the polymer<br>types | 0.35 | 0.84 | 0.8 | 0.001 |
|  | Regardless of the polymer<br>types |  |  | 0.59 | 0.001 |

**Table 3.**
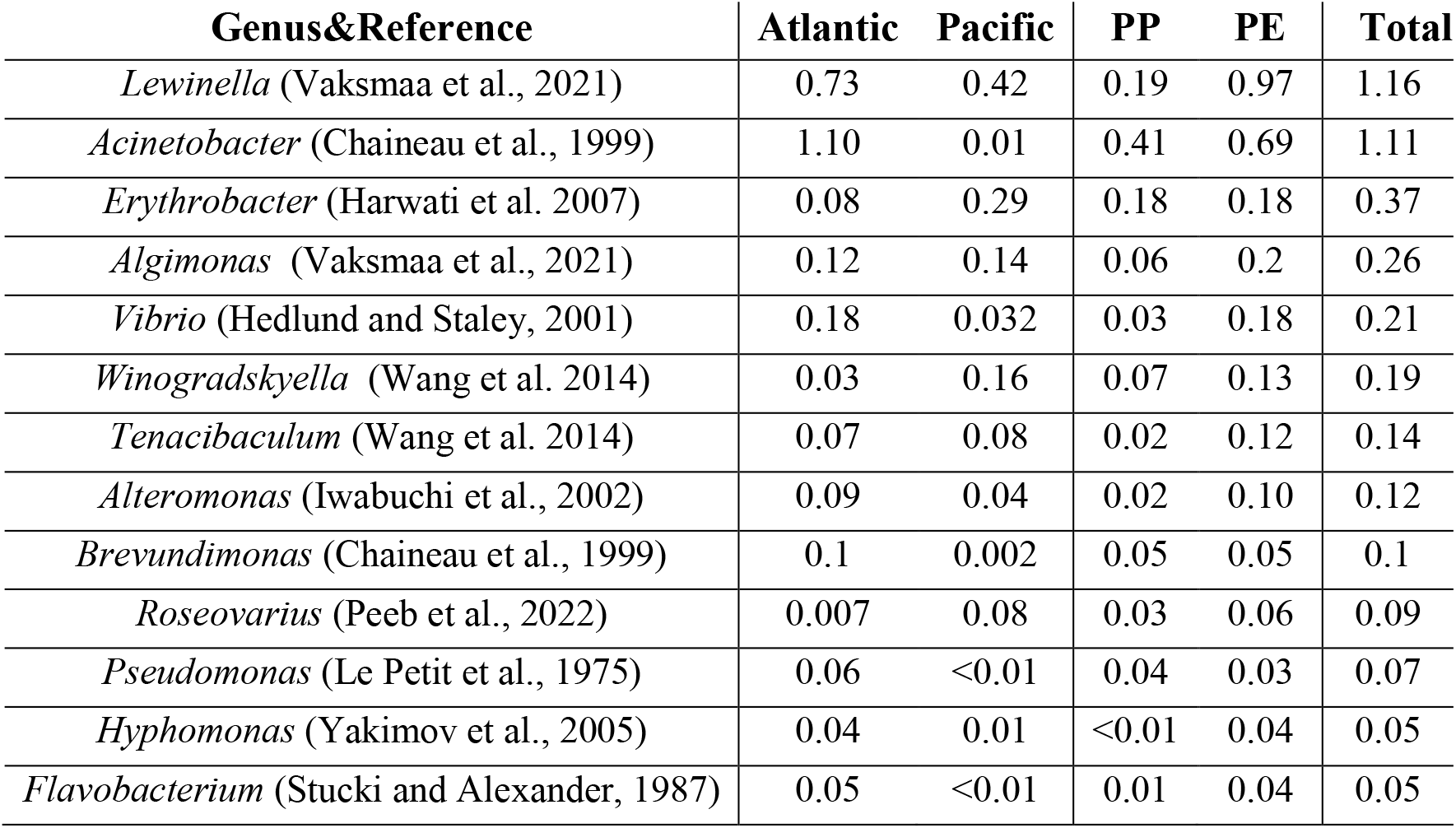
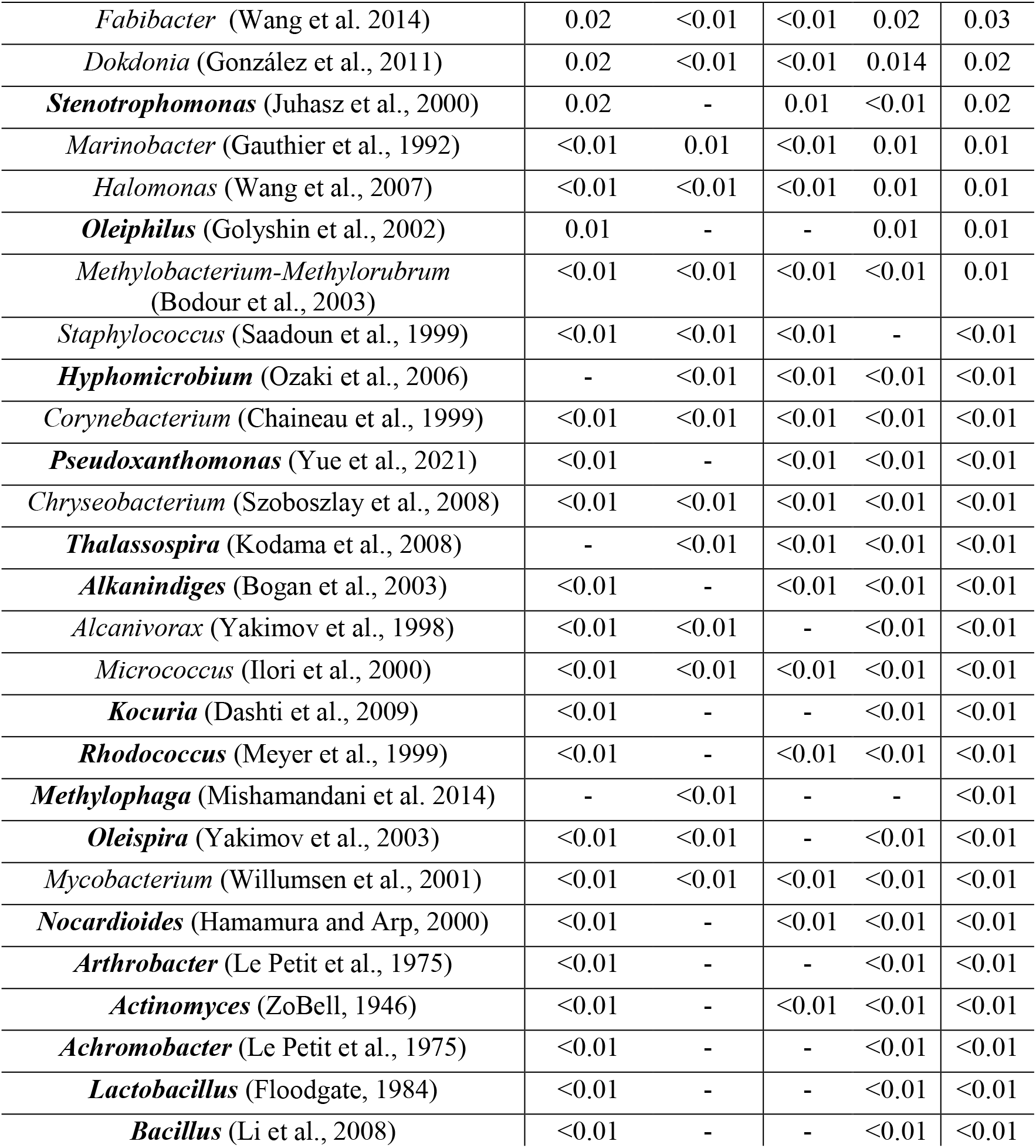
Genera of potential plastic degraders within the studied Plastispheres. Genera in Bold are those detected only in one area, relative abundances are indicated in each ocean and on each plastic type, relative abundances below 0.01 are indicated as <0.01.

### Microbial community composition on the Pacific plastics

After processing, 99.38% of the reads belonged to the domain of Bacteria. Three phyla were most abundant, with almost 91% of the total read count. The other important relative abundant phyla were classified as Planctomycetota, Actinobacteriota and Verrucomicrobiota. They accounted for 6.23% of the total reads. Twenty-seven phyla stemming from Bacteria, Archaea and Eukaryota accounted for 2.76% (each of the 27 recorded below 1% of the reads).

Among the small percentage reads, Archaea (0.62%) showed more diversity in the Pacific than within the Atlantic and were represented by the phyla Thermoplasmatota (0.62%), Nanoarchaeota (0.00058%) and Halobacterota (0.00008%). At the class level, Archaea were represented by Thermoplasmata, Nanoarchaeia and Methanosarcinia. Meanwhile, Eukaryota (0.00018%) displayed less diversity than within the Atlantic. They were represented by one phylum, SAR and one class, Stramenopiles.

Proteobacteria, Cyanobacteria, Bacteroidota and Actinobacteria were the most abundant groups at the phylum level (Figure 4). In addition to the three highest abundant phyla reads, the Pacific Plastisphere recorded Actinobacteria (2.31%), which have been reported as an abundant component of plastic debris communities (Salta et al., 2013; Pinto et al., 2019). Herein, Cyanobacteria and Proteobacteria showed more diversity than in the Atlantic. Gammaproteobacteria were also present, suggesting the maturity of the Pacific biofilms. As such, the Pacific plastics have been drifting for quite some time. At the family level, the figure shows the presence of Hyphomonadaceae and Rhodobacteraceae but not Saprospiracea as in the Atlantic Plastisphere. Instead, Flavobacteraceae, bacteria that prefer surface-attached lifestyles, were present herein.

**Figure 4.**
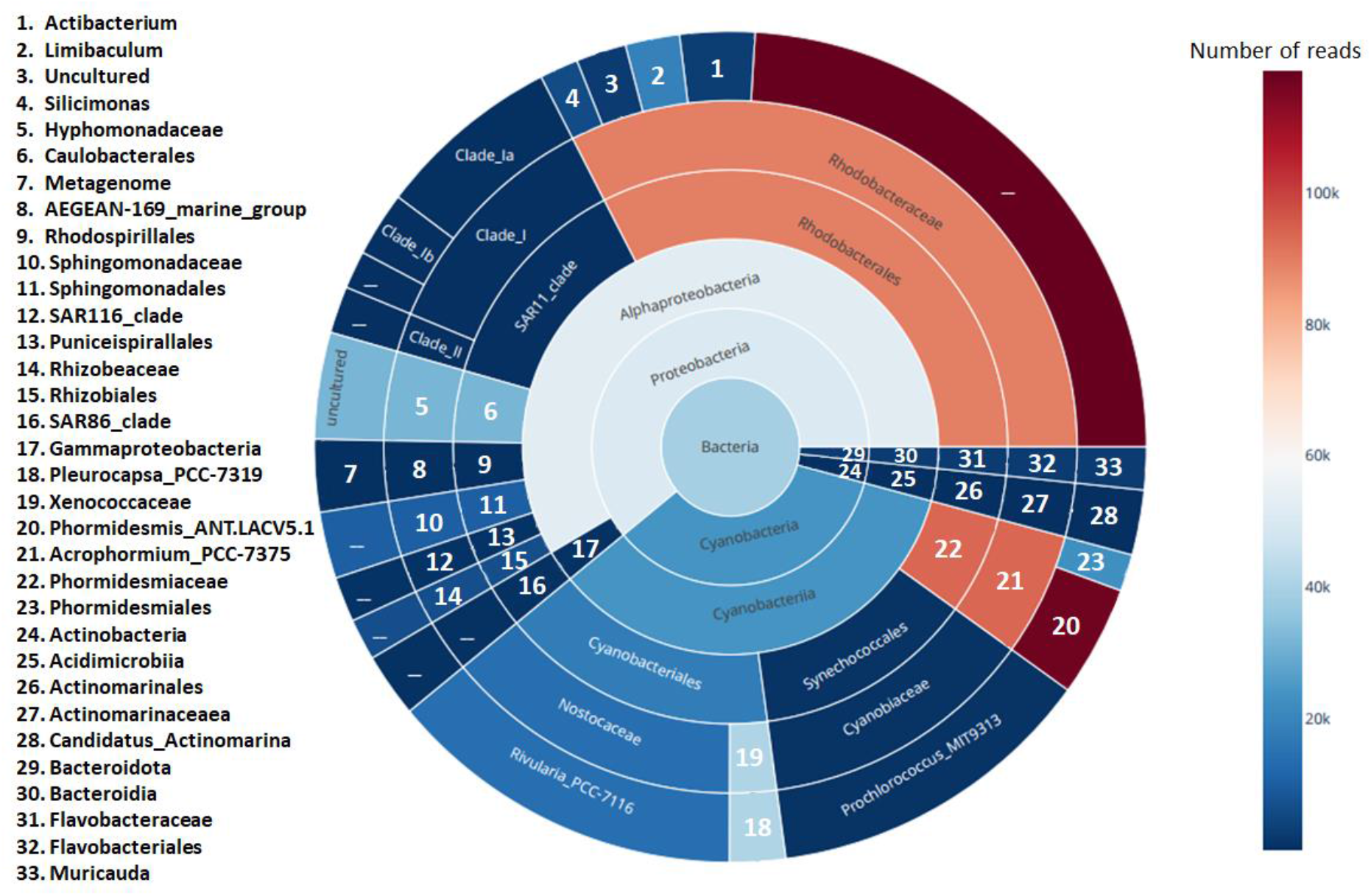
Reads and taxonomic affiliation of recurring communities on the Pacific Plastisphere. Sunburst chart displaying the affiliations of genera that reached values above 20,000 reads. Each plot crown represents one taxonomic level from the Kingdom to the genus. The plot was implemented using ‘plotly.express’ package in Python.

### Microbial community composition in the Pacific water

Pacific water sample analysis was performed to compare microbial communities on Pacific plastic and its surrounding water. Many studies showed that plastic communities differ from surrounding water communities.

From the analysis, Bacteria were the most prominent domain, with 99.62%. Its phyla Proteobacteria, Cyanobacteria and Bacteroidota accounted for more than 91% of the relative abundances. Actinobacteria, Verrucomicrobiota, Planctomycetota and Patescibacteria accounted for 7.11%. The rest (20 phyla), stemming from Bacteria, Archaea and Eukaryota, accounted for 1.83% of the reads.

Archaea in water (0.37%) were represented by the phylum of Thermoplasmatota and the class of Thermoplasmata. Meanwhile, Eukaryota (0.0019%) were represented by the phylum of Amorphea and the class of Obazoa.

Among the occurring phyla between Pacific Plastisphere and Pacific water, Dependentiae (0.005%), PAUC34f (0.002%), Nanoarchaeota (0.0004%, from Archaea), SAR (0.0001%, from Eukaryota), Latescibacterota (0.0001%), Fibrobacterota (0.0001%) and Halobacterota (0.00007%) were found only on Pacific plastic. Amorphea (0.0005%, from Eukaryota) was found only in water. That could probably hint toward the specificity of certain microorganisms for specific substrates.

### Statistical analysis of the microbial community diversity composition of the samples

The statistical analysis of the samples showed a non-significant difference in microbial community diversity within the Atlantic area based on plastic polymer types as well as within the Pacific area. The p-values are greater than 0.05, as shown in Table 2. Indeed, some studies showed that the plastic polymer types have no effect in determining the Plastisphere community composition in mature biofilms (Oberbeckmann & Labrenz, 2020). So, these results confirm the maturity of the biofilms in the Atlantic and Pacific Plastisphere. Also, the p-values displayed (see Table 2) while assessing the diversity between the Pacific plastics and its surrounding water showed no significance. The Pacific Plastisphere was not significantly more or less diverse than the microbial community in the Pacific water. Indeed, Oberbeckmann et al. (2014) suggested that communities at early times in the colonization process are more likely to reveal polymer-specificity, while communities that establish on different polymers should gradually converge over time as the biofilms mature (Harrison et al., 2014).

Meanwhile, the diversity assessment of the Atlantic and Pacific Plastisphere showed significant p-values for phylogenetic measures and beta diversity (see Table 2). So, the communities in the Atlantic Plastisphere are significantly distinct from those in the Pacific Plastisphere. It confirms the results obtained by Amaral-Zettler et al. seven years ago on the same topic when assessing the diversity between Atlantic and Pacific communities. They found the same significance level (p-value = 0.001); distinct grouping based on the oceanic biogeographic zone (Atlantic versus Pacific). Biogeography is incontestably a driver of microbial diversity. Similar results were also obtained by Coons et al., 2021 who found that biogeography influences Plastisphere community structure more than substrate type. Differences in the biofilm community composition are related to different factors.

Some previous studies have targeted temperature as the best predictor of bacterial diversity in surface waters (Ibarbalz et al., 2019). Regarding this study, the plastic particles were collected at the surface of different waters. They could have attracted microbial communities able to evolve at the various water surfaces.

Other studies showed that the substratum physicochemical properties (hydrophobicity, roughness, vulnerability to weather) and the surface chemodynamics (surface conditioning or nutrient enrichment) play a role in microbial diversity (Dang and Lovell, 2016). Besides physicochemical surface properties, it has been shown that the composition of biofilm communities associated with synthetic polymers differed significantly for different ocean basins (Amaral-Zettler et al., 2015) and underlay both seasonal and spatial effects, e.g., in North Sea waters (Oberbeckmann et al., 2014). The waters from the Atlantic and Pacific Oceans seem to have different physicochemical properties, which could have impacted the properties of the plastics we collected, especially since they lasted in the water.

Future studies on the same topic should include environmental parameters to determine the likely drivers of this difference in microbial diversity composition between the Atlantic and Pacific. So, the pH (as it varies between the Atlantic and the Pacific), the dissolved oxygen, the salinity or the surface temperature (as it also varies between both oceans) could be responsible for this difference in microbial diversity between the Atlantic and Pacific oceans.

### Differentially abundant classes between the Atlantic and Pacific Plastisphere

The above statistics showed that there is effectively a significant difference between the Atlantic and the Pacific microbial community diversity. Linear discriminant analysis (LDA) effect size (LEfSe) was used to predict the class level abundances between the Atlantic and the Pacific for their different communities. It revealed 32 differentially abundant classes (LDA log score > ±2) between the Atlantic and the Pacific, as displayed in Figure 5. The dominant classes that made the difference between the Atlantic and the Pacific (see Figure 5) belong to the phyla Proteobacteria, Bacteroidota, Planctomycetota, Bdellovibrionota, Bacilli, Verrucomicrobiota and Thermoplasmatota (from Archaea). SAR and Amorphea (from Eukaryota) were also part of the differentially abundant microorganisms.

**Figure 5.**
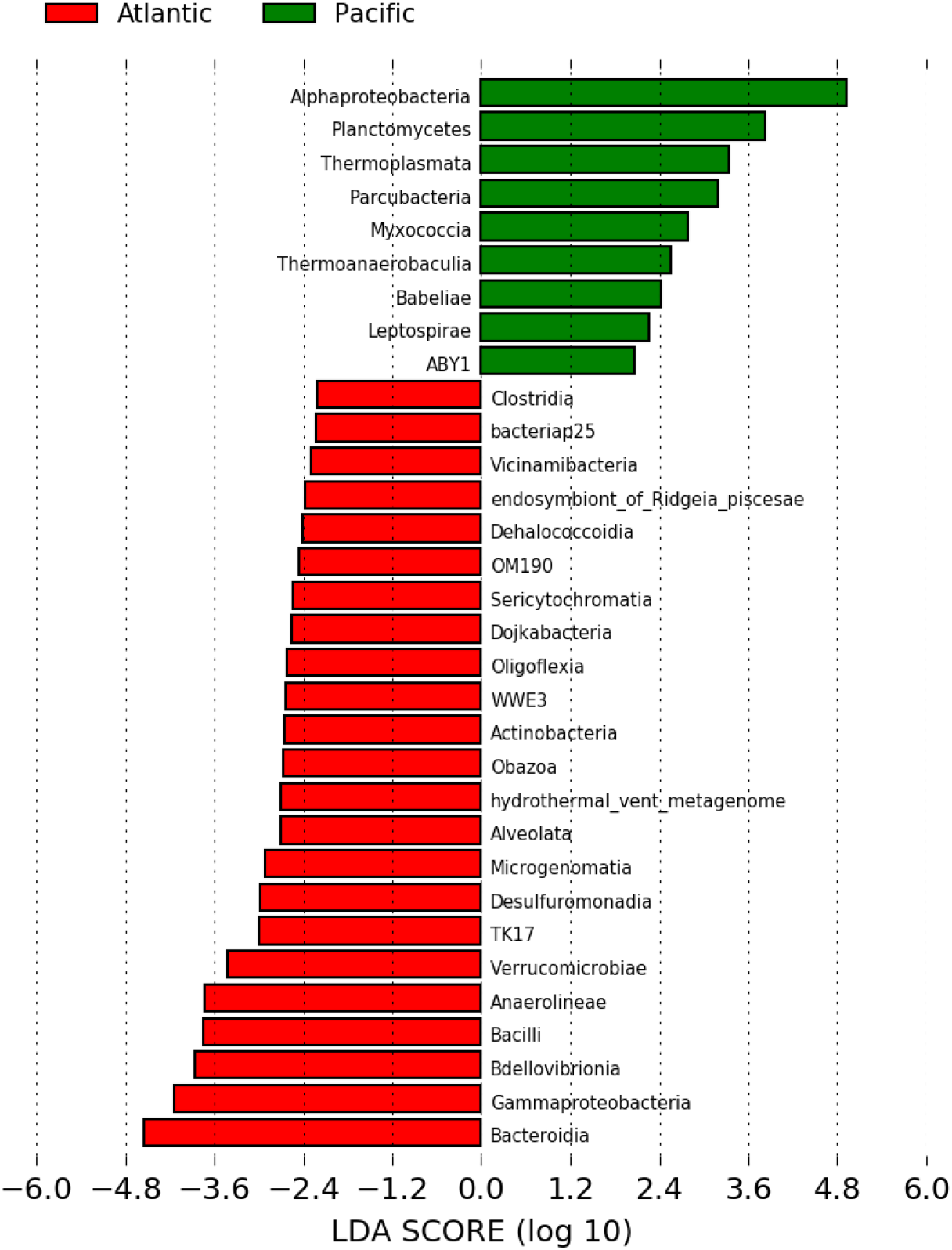
Differentially abundant classes between the Atlantic and the Pacific oceans. Linear discriminant analysis (LDA) effect size (LEfSe) results per ocean. Bar plots depict all classes which had an LDA log score > ±2 between all plastic samples (N = 68) in the Atlantic (n=30) or Pacific (n=38) oceans. The plot was made by utilizing galaxy online (https://huttenhower.sph.harvard.edu/galaxy/). Grouped data were first analyzed using the Kruskal-Wallis test with a significance threshold of 0.05 to determine if the data was differentially distributed between groups.

The Atlantic shows 23 less abundant classes, among which Alveolata and Obazoa are from Eukaryota. In comparison, the Pacific offers nine more abundant classes, among which is the class Thermoplasmata from Archaea. Among these 32 classes, 12 had an LDA score > ±3, including eight from the Atlantic (in ascending order Desulfuromonadia, TK17, Verrucomicrobiae, Anaerolineae, Bacilli, Bdellovibrionia, Gammaproteobacteria) and four from the Pacific (in ascending order Parcubacteria, Thermoplasmata, Planctomycetes, and Alphaproteobacteria). Alphaproteobacteria, Gammaproteobacteria, Bacteroidia had an LDA score > ±4. Thermoplasmata, ABY1 and Desulfovibrionia were unique to the Pacific, while Obazoa, endosymbiont_of_Ridgeia_piscesae, Vicinamibacteria, Alveolata and TK17 were unique to the Atlantic.

### Potential plastic degraders within the studies Plastisphere

The plastic-degrading potential of the Plastisphere community is an ongoing topic (Zettler et al., 2013). Exploring the present Plastisphere, 40 genera previously described to include hydrocarbon-degrading bacteria, as shown in Table 3, were deciphered. These genera represented 4.07% of the relative abundances of the whole Plastisphere and were shared in 4 phyla, five classes, 21 orders and 32 families. Proteobacteria was the most represented, with 22 genera. Actinobacteria came after that with eight genera, Bacteroidota with seven genera and Firmicutes with three genera. Twelve genera were exclusively detected in the Atlantic and three in the Pacific, while 25 were shared between the two oceans.

Our samples were composed of PP and PE. The distribution of PE-degrading microorganisms seems limited, although PP appears to be non-biodegradable. However, it was reported that *Acinetobacter* sp.351 partially degraded lower molecular weight PE oligomers (the genus was found herein: 1.11%) upon dispersion. In contrast, high molecular weight PE could not be impaired (Tsuchii et al., 1980). The biodegradability of PE could be improved by blending it with biodegradable additives, photoinitiators or copolymerization (Griffin, 2007; Hakkarainen & Albertsson, 2004). A blending of PE with additives generally enhances auto-oxidation, reduces the molecular weight of the polymer, and then makes it easier for microorganisms to degrade the low molecular weight materials.

Meanwhile, the possibility of degrading PP with microorganisms has been investigated (Cacciari et al. 1993). In that study, it was shown that aerobic and anaerobic species with different catabolic capabilities could act in close cooperation to degrade polypropylene films. Some *Pseudomonas* (present in this Plastisphere) species were pointed out in the process of polypropylene degradation. Besides that, many species of *Pseudomonas* were indicated to degrade Polyethylene (Zheng et al., 2005), Polyvinyl chloride (Danko et al., 2004), while *Rhodococcus* was shown to degrade Polyethylene (Sivan et al., 2006).

Microbial communities associated with plastic degradation composition and species richness are influenced by spatiotemporal phenomena like habitats/geographical location, ecosystem, and seasonal variation (Kirstein et al., 2019; Pinto et al., 2019). Further, the physiochemical nature of plastics like polyethylene, polypropylene, polystyrene, also regulates this degradation (Pinnell & Turner, 2019). The composition and specificity of microbial assemblage associated with polyethylene (PE) and polystyrene (PS) in the marine aquatic ecosystem (coastal Baltic Sea) are indicated by an abundance of Flavobacteriaceae (*Flavobacterium*), Rhodobacteraceae (*Rhodobactor*), Methylophilaceae (*Methylotenera*), Plactomycetaceae (Planctomyces, *Pirellula*), Hyphomonadaceae (*Hyphomonas*), Planctomycetaceae (*Blastopirellula*), Erythrobacteraceae (*Erythrobacter*), Sphingomonadaceae (*Sphingopyxis*), etc. (Oberbeckmann et al., 2018). Kirstein et al., 2019 found that the microbial community composition associated with various plastics is significantly varying, and it is also changing with the different phases of the plastic degradation process. In our study, the genera, *Flavobacterium* (0.05%), *Hyphomonas* (0.01%) and *Erythrobacter* (0.29%) were precisely found to be associated with PE (0.27%), but also PP (0.2%).

Biotechnologies are leading scientists toward an alternative to mechanical or chemical degradation of plastic waste for a more sustainable end life of plastics. Finding a natural solution to the man-made plastic problem is an urgent issue, but the work keeps going. Once a candidate has revealed plastic degrading potential, the responsible gene must be uncovered and cloned into a host organism (e.g., *Bacillus subtilis;* Austin et al., 2018) to optimize the secreted protein (Wang et al., 2020). Thus, an efficient natural solution against plastic pollution can be found.

## ACKNOWLEDGEMENT

The here presented work was carried out as part of a master thesis project by Dkawlma Tora under the umbrella of the WASCAL (West African Science Service Center in Climate Change and Adapted Land Use) master programm at the Atlantic Technical University Cabo Verde, funded by BMBF (German Federal Ministry of Education and Research) and its partner institutes. We acknowledge the logistical support of Björn Fiedler and Hassan Humeida including ocean-going field work (Floating University 2022) and an extended research stay at GEOMAR. We acknowledge the support of Grace Walls from the Memorial University of Newfoundland, Canada for help during the research cruise POS536. Ute Hentschel and Erik Borchert acknowledge financial support from the BMBF funded project PLASTISEA (GA no.: 031B0867A). Mechthild Schmitt-Jansen and Stefan Lips acknowledge funding of the project P-LEACH (https://www.ufz.de/P-LEACH) from the Helmholtz Association, Innovation Pool of the Research Field Earth and Environment in support of the Research Program “Changing Earth – Sustaining our Future”.

## Supplementary material

**Supplementary table S1**: Samples used in this study with their respective geographic location, plastic type and shannon diversity indice indicated.

