## Supplementary table S1 for "16s rRNA gene sequence analysis of the microbial community on microplastic samples from the North Atlantic and Great Pacific Garbage Patches"

**Supplementary material**

**Supplementary table S1:** Samples used in this study with their respective geographic location, plastic type and shannon diversity indice indicated.

| Sample | Ocean | Plastic type | Shannon diversity |
|---|---|---|---|
| P_1 | Pacific Ocean | LDPE | 6.15 |
| P_2 | Pacific Ocean | HDPE | 6.93 |
| P_3 | Pacific Ocean | PE | 7.48 |
| P_4 | Pacific Ocean | PE | 5.97 |
| P_5 | Pacific Ocean | PE | 6.88 |
| P_6 | Pacific Ocean | PE | 5.36 |
| P_7 | Pacific Ocean | PE | 5.61 |
| P_8 | Pacific Ocean | PE | 6.01 |
| P_9 | Pacific Ocean | PE | 6.62 |
| P_10 | Pacific Ocean | PP | 6.97 |
| P_11 | Pacific Ocean | PE | 6.96 |
| P_12 | Pacific Ocean | PE | 6.69 |
| P_13 | Pacific Ocean | PP | 5.33 |
| P_14 | Pacific Ocean | PE | 6.96 |
| P_15 | Pacific Ocean | PE | 7.26 |
| P_16 | Pacific Ocean | PE | 7.15 |
| P_17 | Pacific Ocean | PE | 6.48 |
| P_18 | Pacific Ocean | PP | 6.72 |
| P_19 | Pacific Ocean | PP | 5.83 |
| P_20 | Pacific Ocean | PP | 5.96 |
| P_21 | Pacific Ocean | HDPE | 5.58 |
| P_22 | Pacific Ocean | LDPE | 7.41 |
| P_23 | Pacific Ocean | PE | 6.41 |
| P_24 | Pacific Ocean | PE | 6.85 |
| P_25 | Pacific Ocean | PE | 7.03 |
| P_26 | Pacific Ocean | PP | 6.92 |
| P_27 | Pacific Ocean | PE | 7.51 |
| P_28 | Pacific Ocean | PE | 5.06 |
| P_29 | Pacific Ocean | PP | 6.46 |
| P_30 | Pacific Ocean | PP | 7.24 |
| P_31 | Pacific Ocean | PE | 6.64 |
| P_32 | Pacific Ocean | PE | 6.57 |
| P_33 | Pacific Ocean | PE | 7.63 |
| P_34 | Pacific Ocean | PE | 5.59 |
| P_35 | Pacific Ocean | PE | 6.77 |
| P_36 | Pacific Ocean | PE | 6.42 |
| P_37 | Pacific Ocean | PP | 6.09 |
| P_38 | Pacific Ocean | PP | 7.33 |
| P_39 | Pacific Ocean | Water | 6.40 |

| | | | |
|---|---|---|---|
| P_40 | Pacific Ocean | Water | 7.34 |
| P_41 | Pacific Ocean | Water | 5.10 |
| P_42 | Pacific Ocean | Water | 6.56 |
| P_43 | Pacific Ocean | Water | 7.52 |
| P_44 | Pacific Ocean | Water | 6.37 |
| P_45 | Pacific Ocean | Water | 5.52 |
| P_46 | Pacific Ocean | Water | 5.48 |
| P_47 | Pacific Ocean | Water | 7.43 |
| P_48 | Pacific Ocean | Water | 7.23 |
| P_49 | Pacific Ocean | Water | 6.78 |
| P_50 | Pacific Ocean | Water | 6.76 |
| P_51 | Pacific Ocean | Water | 7.71 |
| P_52 | Pacific Ocean | Water | 7.30 |
| A_1 | Atlantic Ocean | HDPE | 4.88 |
| A_2 | Atlantic Ocean | HDPE | 6.57 |
| A_3 | Atlantic Ocean | HDPE | 6.31 |
| A_4 | Atlantic Ocean | HDPE | 8.09 |
| A_5 | Atlantic Ocean | HDPE | 5.14 |
| A_6 | Atlantic Ocean | PP | 6.40 |
| A_7 | Atlantic Ocean | HDPE | 6.82 |
| A_8 | Atlantic Ocean | HDPE | 7.26 |
| A_9 | Atlantic Ocean | HDPE | 6.30 |
| A_10 | Atlantic Ocean | HDPE | 7.67 |
| A_11 | Atlantic Ocean | HDPE | 7.60 |
| A_12 | Atlantic Ocean | HDPE | 7.02 |
| A_13 | Atlantic Ocean | HDPE | 7.32 |
| A_14 | Atlantic Ocean | HDPE | 7.30 |
| A_15 | Atlantic Ocean | HDPE | 7.33 |
| A_16 | Atlantic Ocean | HDPE | 7.62 |
| A_17 | Atlantic Ocean | HDPE | 7.13 |
| A_18 | Atlantic Ocean | HDPE | 6.74 |
| A_19 | Atlantic Ocean | PP | 7.82 |
| A_20 | Atlantic Ocean | HDPE | 8.75 |
| A_21 | Atlantic Ocean | PP | 7.76 |
| A_22 | Atlantic Ocean | HDPE | 5.27 |
| A_23 | Atlantic Ocean | PP | 7.48 |
| A_24 | Atlantic Ocean | HDPE | 6.06 |
| A_25 | Atlantic Ocean | PP | 6.51 |
| A_26 | Atlantic Ocean | HDPE | 7.60 |
| A_27 | Atlantic Ocean | HDPE | 5.82 |
| A_28 | Atlantic Ocean | HDPE | 7.01 |
| A_29 | Atlantic Ocean | HDPE | 7.45 |
| A_30 | Atlantic Ocean | HDPE | 7.43 |
